# Complex effects of day and night temperature fluctuations on thermally plastic traits in a seasonal plasticity model

**DOI:** 10.1101/207258

**Authors:** Yara Katia Rodrigues, Erik van Bergen, Filipa Alves, David Duneau, Patrícia Beldade

## Abstract

**Background:** Changes in development in response to seasonally variable environments can produce phenotypes adjusted to fluctuating seasonal conditions and help organisms cope with temporal heterogeneity. In contrast to what happens in natural situations, experimental studies of developmental plasticity typically use environmental factors held constant during development, precluding assessment of potential environment-by-environment interaction effects.

**Results:** We tested effects of circadian fluctuations in temperature on a series of thermally plastic traits in a model of adaptive seasonal plasticity, the butterfly *Bicyclus anynana*. Comparing phenotypes from individuals reared under two types of fluctuations (warmer days with cooler nights, and cooler days with warmer nights) and those reared under a constant temperature of the same daily average allowed us to identify complex patterns of response to day and night temperatures. We found evidence of additive-like effects (for body size), but also different types of “dominance”-type effects where one particular period of the light cycle (for development time) or one particular extreme temperature (for eyespot size) had a relatively larger contribution to phenotype expression. We also gathered evidence against the hypothesis that thermal plasticity in development time drives thermal plasticity in other traits.

**Conclusions:** Combined effects of fluctuating day and night temperatures include additive-like effects as well as different types of environmental-dominance interaction effects. Differences between plastic traits reveal independent responses to temperature, and possible independent assessment of temperature conditions. Our study underscores the importance of understanding how organisms integrate complex environmental information towards a complete understanding of natural phenotypic variation and of the potential impact of environmental change thereon.

## Background

Phenotypic diversity results from complex interactions between organisms and their environments, which happen at different time scales. External environmental conditions contribute to selecting phenotypic variants across generations, but also to generating variation by affecting organismal development. The phenomenon by which environmental conditions affect developmental rates and/or trajectories, leading to the production of distinct phenotypes from the same genotype, is called developmental plasticity [1]. This plasticity is adaptive if the phenotypes generated in response to the environmental conditions experienced during development are better adjusted to the environmental conditions the organisms will experience during adulthood [1,2]. In this manner, plasticity offers a means for organisms to cope with environment heterogeneity, such as that characteristic of alternating seasons. Seasonal polyphenism refers to distinct phenotypes being produced in response to seasonally variable environmental factors, such as temperature and photoperiod [3,4]. Compelling examples include wing development in aphids [4,5], wing pigmentation in butterflies [3,6–8], and diapause in a variety of animals [9,10].

Effects of external environmental factors on development have been amply documented for various traits and species [11,12], as have gene-by-environment (GxE) interactions [13,14]. Efforts to partition genetic effects into additive and interaction components take into account that there are multiple genes and alleles whose individual effects can depend on the genetic context (GxG effects). In contrast, much less attention has been given to potential environment-by-environment (ExE) interactions [15–17]. Traditionally, experimental studies of developmental plasticity have focused on the effects of single environmental factors held constant during the time it takes to complete development. This is in stark contrast with the complexity of natural situations, where multiple and highly dynamic environmental factors can have distinct effects on different genotypes and plastic traits. Towards a more complete account of phenotypic variation, recent studies have started to address phenotypic effects of combinations of different types of cues [18–20]. Less attention has been given to changes in particular environmental factors during development [21,22]. Environmental factors such as temperature fluctuate regularly not only with the yearly seasons, but also with the daily light-dark cycle. Despite the prevalence and importance of circadian fluctuations in ambient temperature, we still lack a clear understanding of the combined effects of day and night temperatures on thermally plastic traits, such as those described for the seasonally plastic butterfly *Bicyclus anynana*.

*B. anynana* has become a valuable experimental model of adaptive developmental plasticity, where we can integrate information about the evolution and ecological significance of plasticity with knowledge about its physiological underpinnings [7,23–25]. In its natural habitat in sub-Saharan Africa, these butterflies typically have two seasonal forms that differ in various traits in association with alternative strategies for avoiding predation and for reproduction. Relative to wet-season form butterflies, dry-season form individuals are larger and delay reproduction until host plants become available for a new generation of larvae [8,26,27]. Dry-season individuals also have less conspicuous wing patterns and their dull brown coloration is thought to provide camouflage against the background of dry leaves, thereby helping resting butterflies escape predators’ attention [8,28,29]. Wet-season butterflies, on the contrary, minimize predator attack by deflecting the attention of predators away from the fragile body, towards their wing margins decorated with conspicuous wing pattern elements called eyespots [30,31]. The main environmental cue determining which form will be produced is the temperature experienced during development [8,32]. Developmental temperature affects the dynamics of ecdysone titres, which, in turn, regulates the response of a suite of plastic traits [24,33]. With only two exceptions [34,35], laboratory studies of *B. anynana* plasticity used temperatures held constant during light and dark hours of the day.

Here, we compared a series of thermally plastic traits between individuals reared under three constant temperatures or under circadian temperature fluctuations with the same daily average as the intermediate constant temperature (Fig. 1a). To probe the effects of the association between temperature and light, we included two regimes with temperature fluctuations: warmer days and cooler nights, as well as the reverse situation. We tested the null hypothesis of no interaction between day and night temperatures by comparing the effect of temperature fluctuations with those of the constant temperature of the same daily average. We also tested the null hypothesis of no association between temperature and light phase by comparing the two types of fluctuations. We found differences between target traits in relation to the combined effects of day and night temperature, including additive and non-additive effects of different kinds. Finally, our data also provide evidence against a previous suggestion that the effect of circadian temperature fluctuations on different thermally plastic traits is a consequence of their direct effects on development time.

**Fig. 1.**
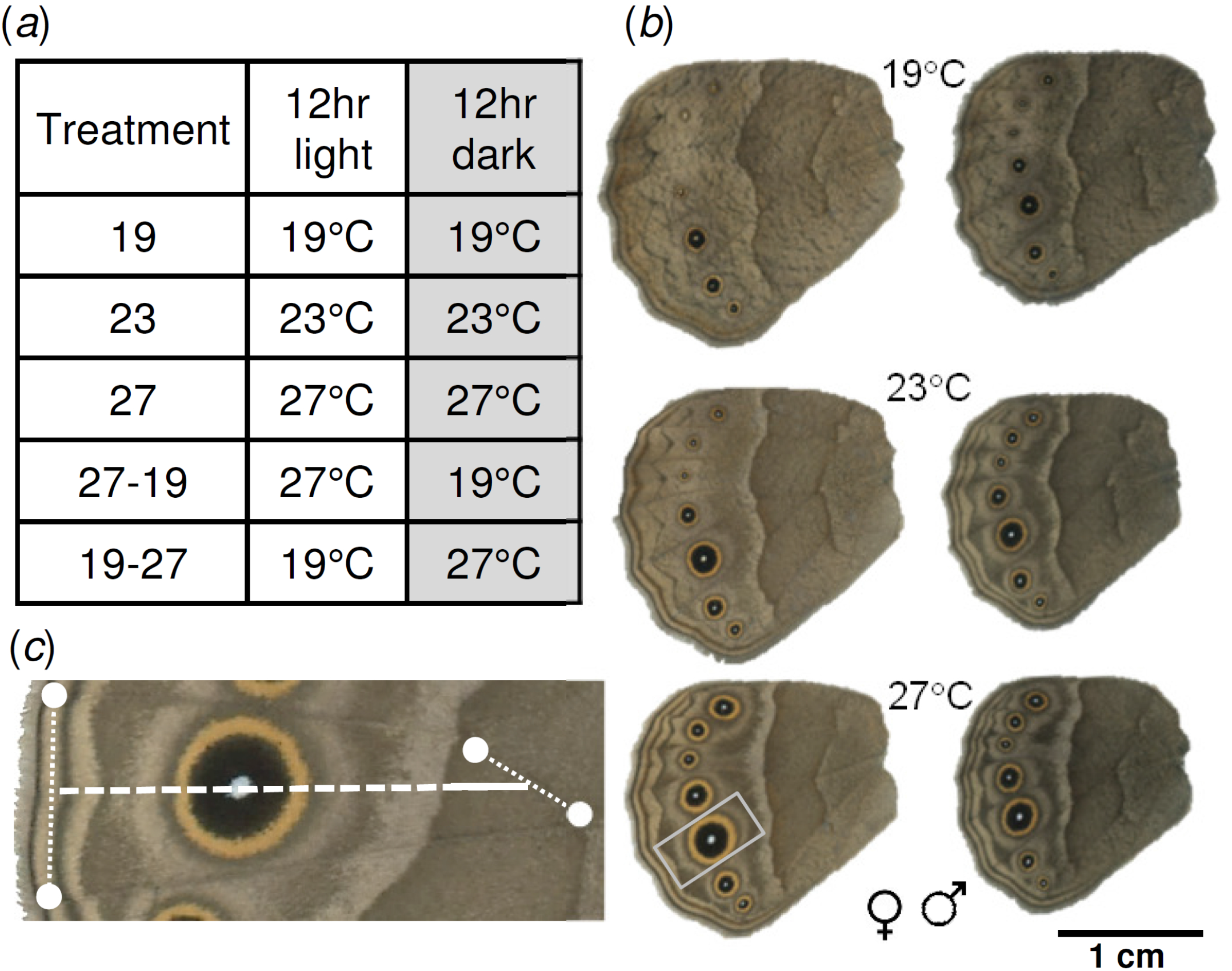
Treatments and wing pigmentation phenotypes. (**a**) Thermal regimes with constant and fluctuating temperatures in association to the light-dark circadian cycle. (**b**) Examples of hindwings (ventral surface) from female and male adults from the different constant temperature treatments. (**c**) Section of a female hindwing (region corresponding to rectangle in panel (**b**) where landmarks (white circles) defined two contiguous transects (white dashed line) passing through the centre of the fifth eyespot. The proximal portion of the transect (solid line) indicates the approximate region used to phenotype the brightness of background.

## Methods

### Butterflies and temperature treatments

We used a captive outbred population of the tropical butterfly *B. anynana* [23] kept in climate-controlled conditions with 65% humidity and 12-12 hrs light-dark cycles (Sanyo MLR-351H or Aralab FITOCLIMA 1000 EH incubators). Caterpillars were fed with young maize plants and adults with sliced banana on wet cotton. To set our experiment, we collected eggs from a large cohort of adults housed at 27°C and allowed them to hatch at the same temperature. Each day for a period of four days, we collected first instar larvae (L1) and randomly assigned them to cages with 22 L1 each that were split into five temperature treatments. Three treatments had constant temperatures: 19°C and 27°C extremes (simulating typical average temperatures of the dry and wet seasons, respectively), and an intermediate of 23°C. Two additional treatments had a daily average temperature of 23°C, but cyclical fluctuations with the light-dark cycle between the two extreme temperatures (Fig. 1a). For each of these five thermal regimes, we had four replicate cohorts in four independent cages. The position of the cohorts within each incubator was changed regularly, and food availability was monitored daily. We checked larval cages daily and transferred pre-pupae into individual cups where they were monitored for pupation and adult eclosion. Adults were allowed to fully stretch their wings before being frozen at −20°C. Wings were dissected and stored at 4°C until phenotypic analysis.

### Quantification of phenotypic traits

We quantified the response to thermal regimes for various thermally-plastic life-history and wing pigmentation traits. We monitored development time by recording the number of days from L1 larvae to pre-pupae, from pre-pupae to pupae, and from pupae to adult, and we calculated total development time by adding those. We measured two proxies of body size: pupal mass and adult wing area. For pupal mass, one-day-old pupae were weighed to the nearest 0.001g (KERN ABS 80-4N scale). For wing area, we scanned the ventral surface of adult hindwings using a colour-calibrated digital scanner (Epson V600) and analysed the resulting images with a set of custom-made interactive Mathematica notebooks (Wolfram Research, Inc., Mathematica, Version 10.2, Champaign, IL, 2015) to measure hindwing area and a series of wing pigmentation traits. For the colour pattern measurements, we first drew two contiguous transects defined by the centre of the fifth eyespot, which is often used to document wing pattern plasticity in this and other species [8,36], and four wing landmarks (on the wing margin and intersection between veins; Fig. 1b-c) in that eyespot’s wing compartment. We marked the limits of each of the colour rings along the transect (central white focus, middle black disc, and external golden ring) to determine ring diameters and calculate the approximate eyespot area (considering it as a circle). The colour of eyespot rings and wing background were quantified using the mean RGB values of the pixels in 3-pixel high rectangles centred on the transect. For the wing background colour, we used the most proximal 50 pixels of the transect, corresponding to a wing region without any defined colour pattern element (Fig. 1c). RGB values were converted to HSB (Hue, Saturation, and Brightness) using the *rgb2hsv* function in R. Background colour was characterized by the brightness value in the HSB colour space; high brightness values corresponding to lighter colours.

### Statistical analyses

We compared phenotypes between temperature treatments, each of which included four replicate cages with up to 22 individuals per cage. All statistical tests were done with R [37], separately for males and females. When appropriate, Normal distribution and homoscedasticity of the residuals were tested with Shapiro-Wilk normality tests and Brush-Pagan tests, respectively. We used a general linear hypotheses test (glht) to test for differences between thermal regimes, followed by Tukey post Hoc pairwise comparisons (alpha=0.05) to ascertain differences between pairs of treatments (package *multcomp* in R).

First, to test for differences in development time, we used a Cox proportional hazards model to determine whether “treatment” influenced the proportions of adult eclosions over time (package *Survival* in R). For the different developmental stages and each sex, we tested the model Coxph *survival (time, eclosion) ~ replicate + treatment*. Second, to test for differences in body size (pupal weight and wing area) and wing pigmentation (eyespot size and wing background colour), we applied a linear model and tested the model: *trait ~ replicate + treatment*. The trait “relative eyespot size” corresponded to the ratio between eyespot area and wing area, for which the assumption of a normal distribution of the residuals was confirmed by a Shapiro test. Finally, to test for the correlation between developmental time and relative eyespot area, we used a correlation test with a Spearman method, with the *p* values corrected and adjusted by the False Discovery Rate (FDR) [38]. The same type of analysis was used to investigate the correlation between developmental time and relative eyespot size using a dataset combining previously published data on development time [39] and on eyespot size [36] in *B. anynana*.

## Results

We tested the effect of circadian temperature fluctuations on different thermally plastic traits: development time (Fig. 2), body size (Fig. 3), and wing pigmentation (Fig. 4). We first compared phenotypes between the three treatments with constant temperatures to assess the direction and strength of thermal plasticity in our *B. anynana* population and experimental conditions. We then compared phenotypes between the three treatments of the same daily average temperature to assess the contribution of day and night temperatures to the phenotype. We found different responses for different traits, including additive and non-additive effects of different types. We finally tested the correlation between development time and eyespot size, using our and another independent dataset (Fig. 5).

**Fig. 2.**
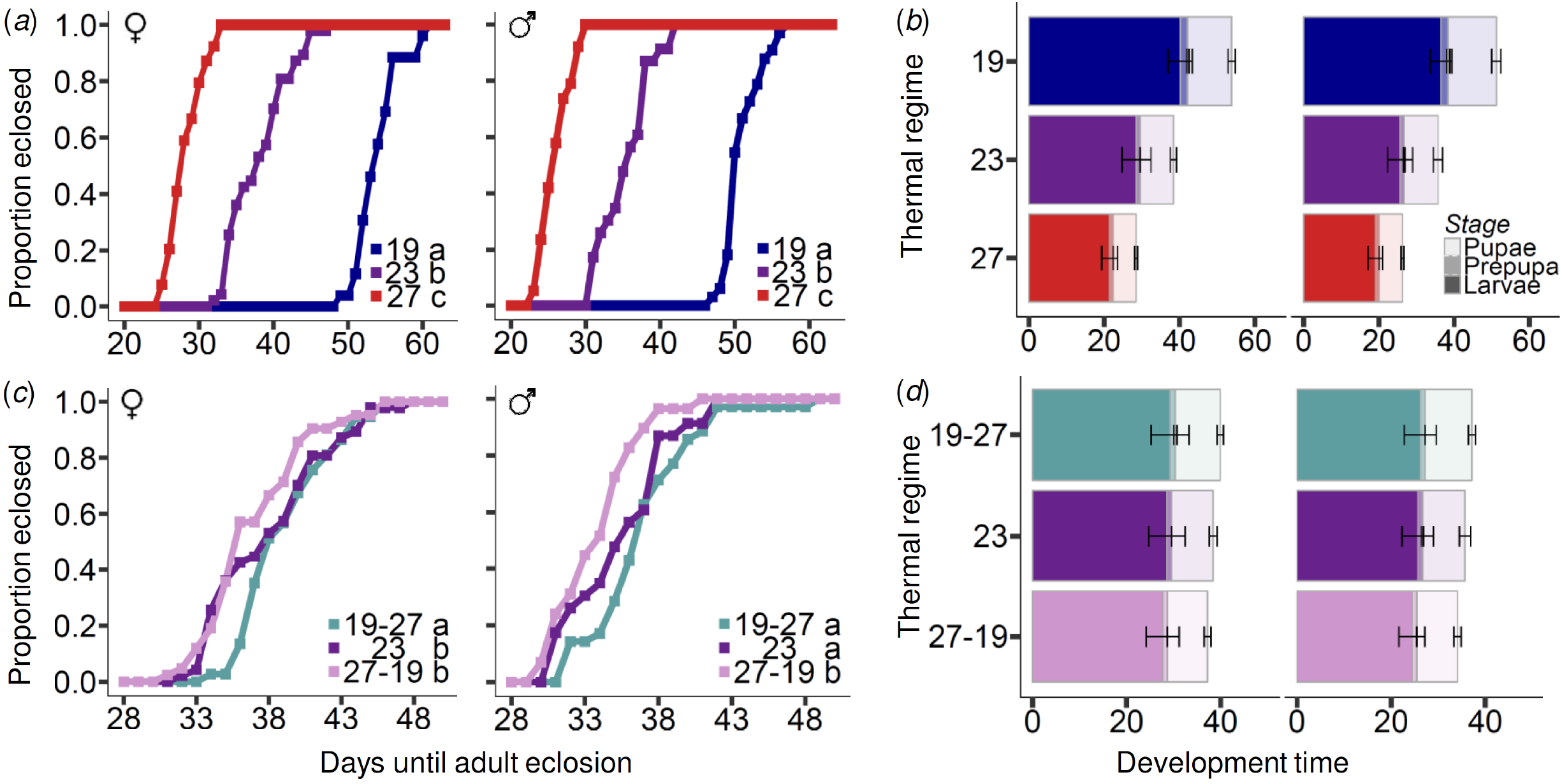
Effects of constant and fluctuating temperatures on development time. Total development time (L1 to adult) and duration of different developmental stages (larvae, pre-pupae, pupae) for females and males developing under constant (**a, b**) or fluctuating (**c, d**) temperatures. Panels (**a**) and (**c**) represent the proportion of adult eclosions since the start of the experiment. Each line corresponds to the individuals of all four replicates for each treatment. There were significant differences (Coxph with *df*=2 and *p*<0.005 in all cases) between constant temperature treatments in (**a**) (*χ*^2^ =245.3 for females, *χ*^2^=202.0 for males), and between the three types of treatments of same daily mean in (**c**) (*χ^2^*=10.6 for females, *χ*^2^=16.8 for males). Letters next to treatment legend illustrate whether pairs of treatments are significantly different (different letters) or not (same letter), *cf*. glth post-hoc test. Panels (**b**) and (**d**) correspond to the duration of different developmental stages. Constant temperature treatments in (**b**) differed in duration of all developmental stages in males (larvae: *χ^2^*=171.2; pre-pupae: *χ^2^*=61.1; pupae: *χ* =170.0) and females (larvae: *χ^2^*=195.9; pre-pupae: *χ* =104.2; pupae: *χ^2^* =203.6; Coxph with df=2 and *p*<0.001 for all cases). Fluctuating temperature treatments in (**d**) differed significantly for the duration of the pupal stage for females (pupae: *χ^2^* =31.0, df=2, *p*<0.001) and males (larvae: *χ* =6.7, *p*<0.03; pupae: *χ* =38.8, *p*<0.001; Coxph with df=2) but none of the other stages (females larvae: *χ^2^* =4.3; pre-pupae: *χ* =4.2 and males pre-pupae: *χ^2^*=0.5, *df= 2*, p>0.05).

**Fig. 3.**
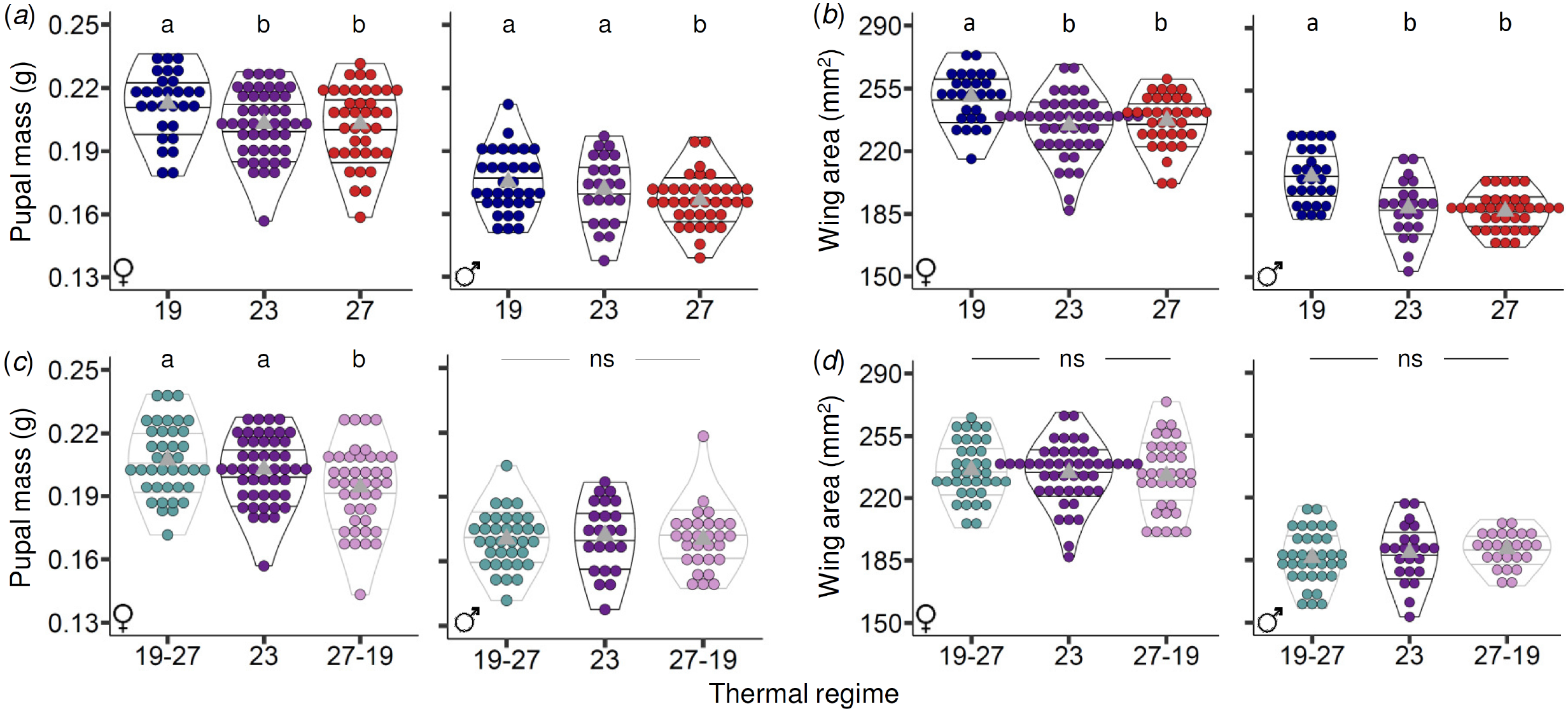
Effects of constant and fluctuating temperatures on body size. Pupal mass and wing area of adult butterflies for females and males developed under constant (**a-b**) and fluctuating (**c-d**) temperatures. Each dot corresponds to one individual (all replicates plotted together) and the red triangles are median values. We found significant differences in pupal mass between constant temperature treatments (**a**) for both females (*F*=4.0, *df*=2, *p*=0.02) and males (*F*=3.1, *df=2, p*=0.04), and between treatments of same daily mean temperature (**c**) for females (*F*=7.5, df=2, *p*<0.001) but not males (*F*=0.1, *df=2, p*=0.91). We found significant differences in adult wing area between constant temperature treatments (**b**) for females (F=16.1, *df=2*, *p*<0.001) and males (F=17.7, *dF=2*, *p*<0.001), but not between treatments of same daily mean temperature (**d**) for females (F=0.6, *df=2*, p=0.56) or males (F=0.9, *df=2*, p=0.40). *ns* refers to non-significant differences between treatments. When there was a significant difference between treatments, letters above treatments illustrate whether they are significantly different (different letters) or not (same letter), *cf*. glth post-hoc test.

**Fig. 4.**
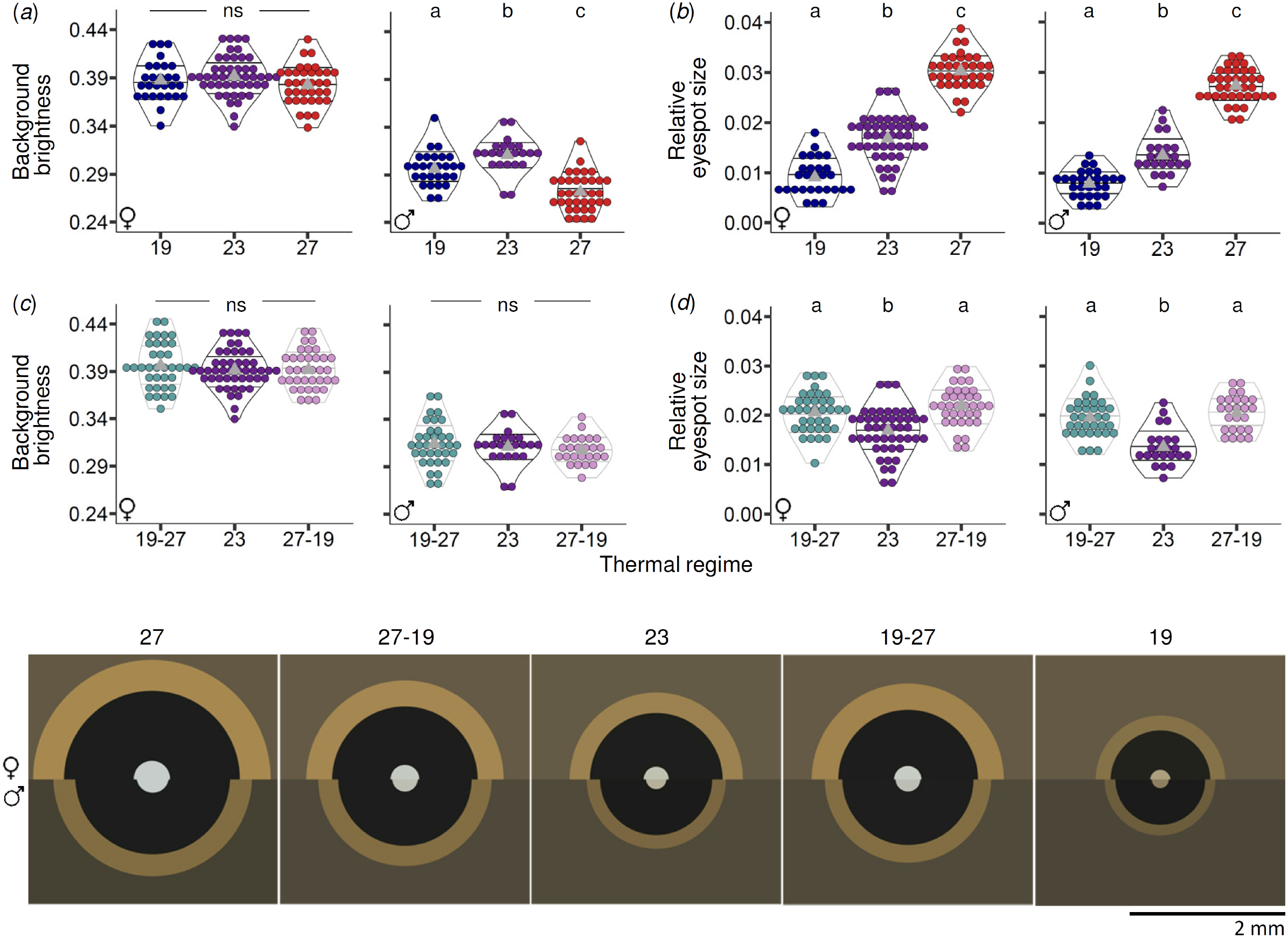
Effects of constant and fluctuating temperatures on wing pigmentation. The background colour and relative eyespot size from females and males developed under constant (a,c) and fluctuating temperatures (b,d). Each dot corresponds to one individual (all replicates plotted together) and the red triangles are median values. We found differences in brightness of wing background colour between constant temperature treatments (**a**) for males (F=33.8, *df=2*, *p*<0.001) but not females (F=2.1, *df=2*, p=0.12), and no significant differences between treatments of same daily mean temperature (**c**) for either sex (F=0.9, *df=2*, p=0.30 for males and females F=0.6, *df=2*, p=0.40). We found significant differences (*ANOVA, df=2*, *p*<0.001 in all cases) in relative eyespot size between constant temperature treatments (**b**) for females (F=223.5) and males (F=315.3), and also between treatments with the same daily mean (**d**) for females (F=14.6) and males (F=25.6). *ns* refers to non-significant differences between treatments. When there was a significant difference between treatments, Letters above treatments illustrate whether they are significantly different (different letters) or not (same letter), *cf*. glth post-hoc test. (**e**) Representation of mean RGB colour for the pixels of the wing background, as well as relative area and colours of eyespot rings from different thermal regimes (see also Additional file 1).

**Fig. 5.**
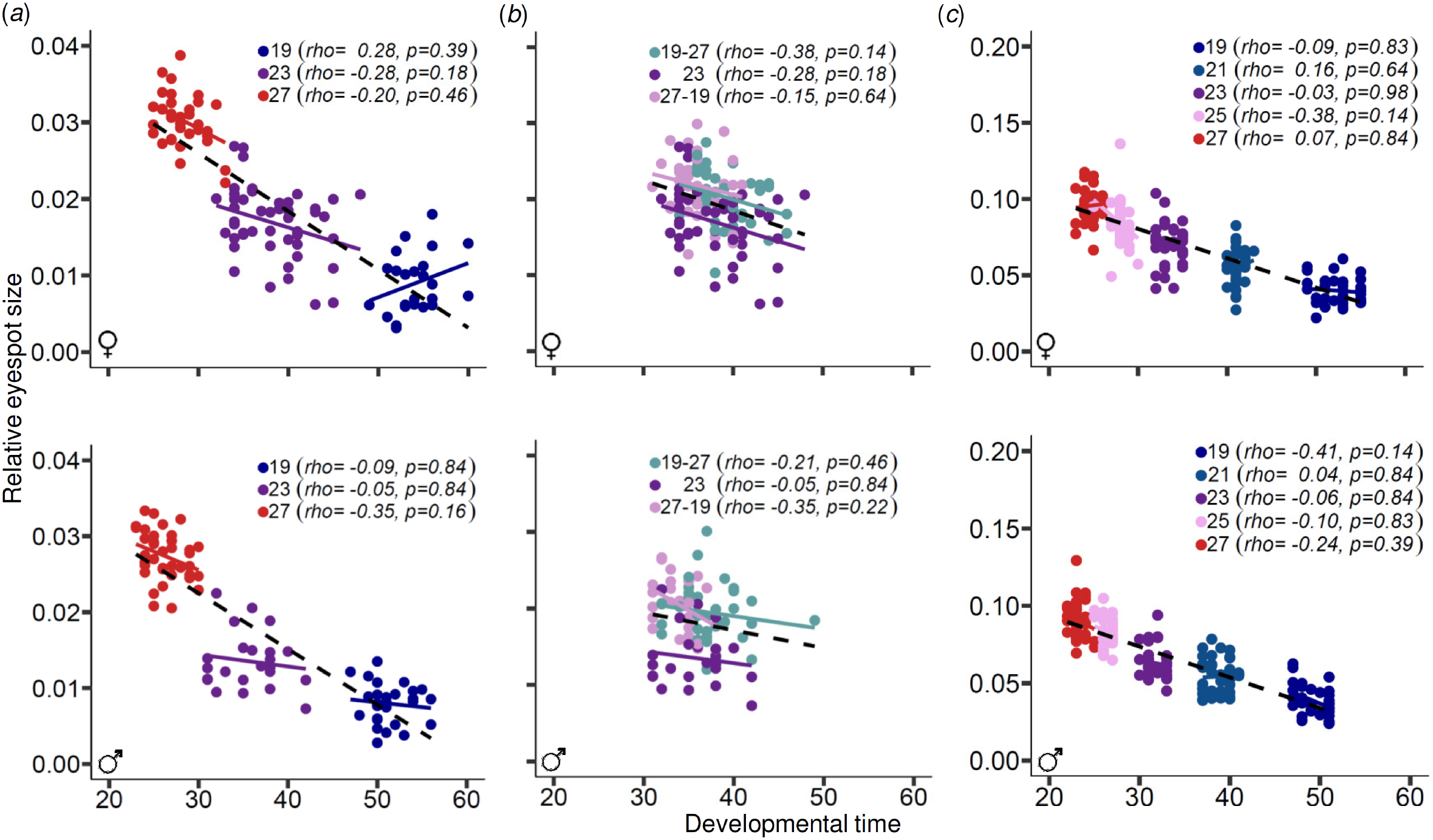
Correlation between relative eyespot size and development time. Relationship between development time and relative eyespot size for females and males from our regimes with constant temperatures (**a**) or with daily mean temperature of 23°C (**b**), as well as data from published work on *B. anynana* using constant temperatures (**c**). Each dot corresponds to one individual and all replicates are plotted together, separately for females and males. Lines correspond to the best fit line: same colour as dots for relationships for data points of the different thermal regime, and black for relationship across all data points. Spearman’s rank correlation coefficient (Spearman rho) test showed a significant negative correlation when data points from all treatments were considered together (black line): (**a**) rho=-0.85 for females and rho=-0.88 for males (*p*<0.001 for both), (**b**) rho=-0.19 (p=0.08) for males and rho=-0.28 for females (p=0.001), (**c**) rho=-0.87 for males and rho=-0.85 females (*p*<0.001 for both). For the correlations within treatments, *rho* and corresponding p-values are given in the figure.

### Different contributions of day and night temperatures to development time

We confirmed thermal plasticity in *B. anynana* development time in our study population: individuals reared at lower temperatures took longer to reach adulthood than individuals reared at higher temperatures (Fig. 2a). For both males and females, temperature affected the duration of all developmental stages monitored; individuals from warmer temperatures had shorter larval, pre-pupal, and pupal stages (Fig. 2b).

We also found differences in development time between the three treatments with a daily average temperature of 23°C (Fig. 2c). For both males and females, development was faster for individuals that spent the day at 27°C and the night at 19°C (27-19 treatment) compared to individuals that spent the day at 19°C and the night at 27°C (19-27 treatment). The duration of the pupal stage differed between treatments, but not the duration of the larval and of the pre-pupal stages (Fig. 2d). The difference between our two treatments with fluctuating temperatures revealed that the effect of day and night temperature on development time is not additive and that the temperature experienced during the light phase had a larger impact on total development time. Individuals reared with a day temperature of 27°C demonstrated a shift in development time towards individuals reared at constant 27°C, while the development time of individuals reared with a day temperature of 19°C shifted towards those reared at a constant temperature of 19°C. The response for individuals reared at a constant temperature of 23°C relative to the two fluctuations of the same daily mean (27-19 and 19-27) differed between males and females (Fig. 2c-d).

### No difference between fluctuations and constant daily temperature for body size

For both proxies of body size we quantified, pupal mass (Fig. 3a) and adult wing area (Fig. 3b), we confirmed known thermal plasticity patterns, with lower temperatures yielding larger individuals. For both sexes, individuals reared at 19°C were significantly larger than individuals reared at 23°C or 27°C, which did not differ significantly between each other.

Overall, we found no significant differences between individuals reared under constant versus fluctuating temperatures of the same daily average (Fig. 3c-d). This corresponds to an additivelike effect of day and night temperatures on body size. The only exception was female pupal mass, where we found a significant difference between the two fluctuating temperature treatments. Like for development time, the temperature experienced during the light phase had a stronger effect on the phenotype (Fig. 3c).

### Different contributions of cool and warm temperatures to eyespot size but not wing background brightness

We investigated two aspects of wing pigmentation (Fig. 4): relative eyespot size, which is a trait well known to be thermally plastic, and wing background colour, which had not been quantified before despite suggestions of it also varying between seasonal forms. As had been noticed but not formally quantified before, females are lighter than males and wing background colour depends on rearing temperature, but only for males (Fig. 4a). Plasticity for eyespot size was much stronger, with significant differences between all three constant temperature treatments (Fig. 4b). This is in line with the well described thermal plasticity for *B. anynana* eyespot size, with larger eyespots in animals reared at warmer temperatures, for both males and females.

Regarding the comparison between constant and fluctuating temperatures of daily average of 23°C, we found similar results for males and females: no differences for wing darkness (Fig. 4c) and clear differences for eyespot size (Fig. 4d). Individuals reared at either of the two fluctuating temperature regimes had larger eyespots than those reared at the constant temperature of 23°C, and were not significantly different from each other. Results for overall eyespot area were consistent with those for the area of individual eyespot rings (central white focus, middle black disc, and external golden ring), which also showed sex and temperature differences in actual colours (Fig. 4e and Additional file 1).

### Correlation between eyespot size and development time between but not within temperature treatments

It had been previously suggested that thermal plasticity in traits such as eyespot size, rather than a direct response to temperature, is a correlated response to temperature-induced changes in development time [29,35,40]. This is not consistent with our results which show that individuals reared at 19-27 developed slower than those from 27-19, but both had larger eyespots than individuals reared at 23°C. We thus went on to investigate the correlation between development time and eyespot size, both across and within temperature treatments. We used our dataset with five thermal regimes, as well as an additional independent dataset put together from published work that includes two extra constant temperature treatments [36,39] (Fig. 5).

Across constant temperature treatments, with largely non-overlapping development times, we found an overall strong negative correlation between development time and eyespot size, for both females and males (Fig. 5a-b). However, within temperature treatments, there were no correlations between development times and relative eyespot sizes that were statistically significantly different from zero. This result was confirmed using an independent dataset from previously published studies that included more intermediate temperature treatments (Fig. 5c).

## Discussion

We investigated the effects of combinations of day and night temperatures on a series of thermally plastic traits in *B. anynana* butterflies. We confirmed and quantified thermal plasticity in our experimental population and conditions for development time, body size, and eyespot size, and documented thermal plasticity in wing background colour. Butterflies reared under warmer temperatures generally had faster development, smaller bodies and larger eyespots, matching the seasonal polyphenism described for the species [6,7,24,25]. To assess potential interaction effects of day and night temperatures, we then compared phenotypes from individuals reared under two types of circadian temperature fluctuations and under a constant temperature of the same daily average.

### Combined effects of day and night temperature on thermally plastic traits

If day and night temperatures contributed equally to phenotype expression, i.e. if their effects were purely “additive”, to borrow from the terminology used to partition genetic effects, we should have no difference between the two types of fluctuations (our 27-19 and 19-27 regimes), and also no difference between those and the treatment with constant temperature of the same daily average (23°C). We found evidence of such additive effects (for body size; Fig. 3), but also of “dominance”-type effects where one particular period of the light cycle (for development time; Fig. 2) or one particular extreme temperature (for eyespot size; Fig. 4) had a relatively larger contribution to phenotype.

Two previous studies had addressed the effect of circadian temperature fluctuations on *B. anynana* but used only warmer days than nights [34,35]. While this is the more ecologically-relevant regime, in isolation, it did not allow identification of interactions between temperature and light phase. Like in those previous studies, we found that butterflies that developed under cooler nights, relative to those that developed under constant temperature of the same daily average (*i.e.* 27-19 versus 23 regimes), had faster development (Fig. 2) and larger eyespots (Fig. 4), but mostly did not differ for the other traits under study (except female pupal mass; Fig. 3). Moreover, we showed that cooler nights speeded up development largely by shortening the duration of the post-feeding pupal phase (Fig. 2b). The explanation previously proposed to account for such effects on total development time was that *B. anynana* caterpillars eat mostly during the dark hours and assimilate those resources during light hours [34]. Cooler nights can presumably sustain higher activity levels and higher feeding rates, while warmer days might allow higher assimilation efficiency. Either or both of these could result in faster development. Relationships between temperature and food ingestion efficiency [41], as well as between thermal stress and depletion of energy reserves [42] have been reported for different arthropods. Another factor that may explain different contributions of day and night temperatures relates to the open question of how often and when developing organisms “acquire information” about external environment [43,44]. Within windows of sensitivity during development (e.g. [24,32,45–47]), it remains unclear whether organisms assess environmental conditions continuously or at discrete time points. The “dominance” effect of the conditions experienced during the light hours on development time could reflect discrete sampling of the environment mainly occurring during that period of the day.

Our comparison between the two types of fluctuations allowed us to gain new insight on the combined effects of day and night temperatures on development time and eyespot size, two iconic examples of *B. anynana* seasonal plasticity. We found distinct types of interaction effects for the two traits. For development time, we found that the temperature experienced during the day had a stronger effect than the temperature experienced during the night. In the same way that individuals develop faster at 27°C relative to 19°C (Fig. 2a), individuals from 27-19 (i.e. day spent at 27°C) developed faster than those from 19-27 (day at 19°C) (Fig. 2c). On the other hand, we found that any period of the light-dark cycle spent at a warmer temperature lead to increased eyespot size, as is characteristic of development at warmer temperatures (Fig. 4a). Individuals developing under both types of temperature fluctuations had larger eyespots than those from the constant temperature of the same daily average (Fig. 4b).

### Independent effects of temperature on different traits that make up a plasticity syndrome

Typically, seasonal forms differ in a suite of traits that respond to seasonably variable environmental conditions [36,39]. In the case of *B. anynana*, this thermal plasticity “syndrome” includes the traits monitored here, as well as others such as starvation resistance, longevity, and reproductive investment [23,39,48]. Supported also by laboratory data on correlated responses to selection on development time [40], it had been suggested that temperature affects development time directly, and it is the ensuing changes in development time that lead to changes in other thermally plastic traits [29,35,40].

Indeed, butterflies developing at lower temperatures take longer to complete development and have smaller eyespots than those developing at warmer temperatures. However, within a thermal regime, the variation in development time between individuals, which can be of several days, did not correlate with eyespot size. This apparent case of the Simpson’s paradox or Yule–Simpson effect [49] was true for our dataset and for data from other independent studies (Fig. 5). These results suggest that temperature-induced changes in development time cannot account for temperature-induced changes in all other thermally plastic traits, and certainly not for changes in eyespot size seen under fluctuating temperatures, and argue for a more direct and trait-specific effect of temperature. Additional support for this comes from the different shapes of reaction norms of traits belonging to the thermal plasticity syndrome, and from the fact that manipulations of the ecdysone titres known to mediate this plasticity have trait-specific effects [33,50]. The trait-specific responses to environmental conditions are probably related to trait-specific windows of environmental sensitivity during development [32,46,47]

### Effects of circadian temperature fluctuations on development and evolution

Experimental studies on different animal and plant systems have documented effects of day-night temperature fluctuations on both development (i.e. phenotype expression) and evolution (i.e. phenotype filtering by natural selection). Examples of the former include effects of day versus night temperature on the regulation of flowering time [51,52], and effects of circadian temperature fluctuations on various fitness related traits [53–57]. The close association between effects of light and of temperature on biological processes is further revealed in the overlap in sensing mechanisms for the two cues (e.g. role of phytochromes as thermosensors in *Arabidopsis* [58,59], or cryptochrome in *Drosophila* [54]), and also the observation that temperature, and not only light, can reset the circadian clock [60]. At the scale of intergeneration effects, evolution under different thermal regimes in natural and experimental populations has documented effects of circadian temperature fluctuations on a variety of phenotypic traits including body size [56,57], as well as on allele frequencies [61].

Unlike most experimental studies of thermal plasticity, we addressed the effects of development under temperatures not held constant. Specifically, we studied potential interaction effects of fluctuating day and night temperatures on a suite of thermally plastic traits. Circadian fluctuating temperatures are undoubtedly closer to reality than constant temperatures, the same way that colder nights are closer to reality than warmer nights. This is the scenario under which organisms have evolved in natural populations, but is rarely the scenario under which animals are maintained or studied in the laboratory (but see e.g. [62]). In fact, even though exposure to temperature change can be used as a form of stress (e.g. [57,63]), it is possible that thermal constancy also constitutes a type of stress [64]. Whether temperature change is or not perceived as a stress capable of triggering stress responses likely depends on how abrupt (rather than gradual), how large, and how recurrent the change is [65]. Many studies of thermal stress use rather short exposures to extreme temperatures (e.g. [11,66]). Such temperature changes can affect different aspects of the organisms’ biology, including developmental robustness [67] and various fitness-related traits [42]. It is unclear if the temperature changes, which typically fluctuate with the day and night cycle, can illicit any of the same type of physiological responses. It is even less well known how organisms integrate complex environmental information, such as that where multiple environmental factors change during the time it takes to complete development, and still produce coherent phenotypes for the various plastic traits [67,68]. Especially when environmental challenge includes a mismatch in what are the usual combinations of environmental factors, environment-by-environment effects are likely to have an important impact on how organisms deal with such challenge.

## Conclusions

We found evidence for different types of combined effects for day- and night-time temperatures on a suite of thermally plastic traits associated with distinct seasonal strategies for survival and reproduction in *B. anynana* butterflies. While, for some traits, day and night temperatures seem to have largely additive effects on phenotype expression, we also identified different types of non-additive effects. These included environmental dominance-like effects where one particular period of the circadian cycle or one particular extreme temperature had a relatively larger contribution to phenotype. Differences between traits reveal their independence in the response to temperature, which might relate to trait-specific windows of environmental sensitivity and/or trait-specific assessment of environmental conditions. Our study underscores the importance of understanding how organisms integrate complex environmental information towards a complete understanding of natural phenotypic variation and of the potential impact of environmental change thereon

## List of abbreviations

GxE: gene-by-environment interactions
GxG: gene-by-gene interactions
ExE: environment-by-environment interactions
L1: first instar larvae

## Declarations

### Ethics approval and consent to participate

Not applicable.

### Consent for publication

Not applicable.

### Availability of data and material

Raw data are available in Additional file 2.

## Authors’ contributions

Y.K.R. and P.B. conceived and designed the study; Y.K.R. performed the experiments and collected the data; F.A. developed a set of interactive Mathematica notebooks to collect wing phenotypic data; E.v.B. provided data on the extra constant thermal regimes and helped collect wing colour data; Y.K.R., E.v.B., and D.D. performed the statistical analyses; Y.K.R. and P.B. drafted the manuscript, with input from E.v.B. and D.D. All authors gave final approval for publication.

## Acknowledgements

We thank Elvira Lafuente for help with data analyses, Vicencio Oostra and Maaike de Jong for access to their published data, and Carolina Peralta and Pedro Castanheira for help with animal husbandry.

## Additional files

Additional file 1: Effects of constant and fluctuating temperatures on eyespot colour rings.

Additional file 2: Dataset used in this study.

